# Co-transcriptional splicing of the *Drosophila* Y-loop A transcript is organized in chromatin hubs and shows variable delay which increases with spermatocyte maturation

**DOI:** 10.64898/2026.11.11.766038

**Authors:** Stefan A. Koestler, Robert White

## Abstract

Splicing of the primary transcript is a major step to produce the mature mRNA. While the biochemistry of the splicing mechanism has been elucidated in detail, the dynamics and structural arrangements inside the nucleus are not clear. Biochemical and genomic studies of splicing and transcription most often lose the information about individual states and localization from the sub-nuclear to the tissue level by averaging over populations and homogenization of cells and tissues. Imaging approaches, which would overcome these problems, have been hampered by the short length of most genes, preventing discrimination between different regions like introns and exons along the transcript due to limited resolution.

To overcome these limitations, we apply high-resolution microscopy with Airyscan joint-deconvolution to the megabase transcription units of *Drosophila* Y-loops. Along these extended chromatin fibers, we can directly observe the linear arrangement and local splicing events of introns and exons as well as the corresponding 3D chromatin structure. We compare these events between primary spermatocytes at different stages of their development within whole-mount testes.

We show that co-transcriptional splicing of the Y-loop A transcript exhibits delayed splicing. The extent of this delay differs between transcription clusters along the same chromatin fiber within a nucleus and the average splicing delay increases with spermatocyte maturation. This shows that co-ordination of splicing and transcription is regulated locally at the site of transcription and by developmental state. We find that the concentration of the transcriptional activator Eaf is inversely correlated with spermatocyte maturation thus suggesting that the decreased frequency of transcription pause/pause release may be responsible for the splicing delay. Consistently, we find hubs containing the spliceosome with elongating and paused RNA-polymerase together with Eaf in chromatin bends. From their different arrangements we derive a model for splicing hub biogenesis.

## Introduction

Transcription of genetic information into mRNA is an essential step and a key control point of cellular protein production. The central player of mRNA transcription is DNA dependent RNA polymerase II (RPol), whose activity can be regulated by phosphorylation of repeat heptad sequences at Ser2 or Ser5 (RPol-PSer2/5) of its C-terminal domain (CTD), thereby acting as a switch between transcription pause and elongation. Phosphorylation of RPol-Ser2 is carried out by Cdk9 of the transcription factor complex p-TEFb which in its most active form is part of the Super Elongation Complex (SEC) (Cramer, 2019). Transcription by RPol is interconnected with various stages of pre-mRNA processing, such as capping, splicing, and polyadenylation (Bentley, 1999).

Splicing is performed by the spliceosome, a complex composed of various proteins and small nuclear RNAs. The assembly of the spliceosome is a multistep process involving the interaction of U1, U2, U4/U6, and U5 snRNPs, along with non-snRNP splicing factors, with the pre-mRNA and each other to define the intron splice sites and the branchpoint (Chathoth et al., 2014). Around 80% of introns in S. cerevisiae, *Drosophila* and Homo sapiens are spliced out co-transcriptionally (Neugebauer, 2019), although it was recently suggested that this number is an overestimation due to limitations of the applied methods (Coté et al., 2024). Both transcription and splicing likely take place in phase separated condensates.

These condensates are dynamic and transition from transcription initiation to elongation, and splicing leads to formation of ‘gene-body condensates’ or splicing speckles (Cramer, 2019; Guo et al., 2019; Pancholi et al., 2021). The SEC plays a major role in this transition by driving the phosphorylation of RPol-PSer2 (Luo et al., 2012a).

While splicing can take place immediately after splice site transcription, it can also be delayed for many kB (Shine et al., 2024; Sousa-Luís et al., 2021). Delayed splicing might have a function, e.g. it has been suggested that it increases chromatin stiffness to form loops (Leidescher et al., 2022), or to allow alternative splicing (Shenasa and Bentley, 2023). It might also be a requirement for or a consequence of fast transcription since splicing has been associated with paused RPol-PSer5, potentially allowing time for quality control at a splicing checkpoint (Alexander et al., 2010). This is in line with the suggestion that in highly expressed genes delayed splicing is more prominent (Ullrich et al., 2025). On the other hand, recently the evidence for pausing at splice sites has been questioned (Shine et al., 2024).

To investigate the architecture of splicing within intact nuclei we applied high-resolution Airyscan joint-deconvolution microscopy to the *Drosophila* Y-loop model system. *Drosophila* Y-loops are tens of µm long chromatin loops that extend from the Y-chromosome into the nucleoplasm of primary spermatocytes and can easily be observed with the light microscope (Bonaccorsi et al., 1988). That transcription and splicing on Y-loops is a representative model system is corroborated by the finding of massive chromatin loops in a range of organisms including humans (Leidescher et al., 2022). Like most genes, the *Drosophila* Y-loops exhibit co-transcriptional splicing (Fingerhut et al., 2024).

To assess the coordination of splicing and transcription along a specific gene within the 3D chromatin environment at a single cell level we applied RNA-FISH with specific exon/intron combinations on the Y-loop A gene *kl-5* of *Drosophila* spermatocytes within whole-mount testes. In testes, spermatozoa are continuously generated from germ line stem cells via spermatogonia and spermatocytes. Primary spermatocytes go through a long G2 phase of several days during which spermatocyte specific transcription is initiated, their nuclei expand 25-fold, and 3 large chromatin regions loop out from the condensed Y-chromosome into the nucleoplasm. Each of the three Y-loops encodes a single gene, each several megabases long, and exons are separated by long introns so that their linear arrangement can be easily studied by RNA-FISH (Bonaccorsi et al., 1988; Fingerhut et al., 2019; Redhouse et al., 2011). The *kl-5* primary transcript encoding the axonemal dynein heavy chain is 3.2 Mb long (Gatti and Pimpinelli, 1983) and this generates an mRNA of 13.6 kb (Flybase (Öztürk-Çolak et al., 2024)). With Airyscan microscopy and joint-deconvolution a resolution of 90 nm can be achieved (Prazeres da Costa et al., 2021). This approach allowed us to determine the position of splicing events along Y-loop A, the arrangement of splicing and transcription factors, and the corresponding chromatin structure within a nucleus. The use of whole-mount testes enabled us to compare natural variations in splicing organization between spermatocytes of different maturity, since the cells are arranged linearly inside the testis according to their consecutive developmental stages. In contrast to bulk and single cell sequencing methods as well as most previous microscopy-based approaches including expansion microscopy (Coté et al., 2024), our approach allowed us to directly visualize and compare local splicing clusters along the same chromatin fiber and between nuclei of cells at different developmental stages within the same tissue.

We show that co-transcriptional splicing of the *kl-5* gene transcript exhibits delayed splicing and that this delay, of up to several hundred kb after the splice site, differs between clusters along the same chromatin fiber and increases with spermatocyte maturation. We complement these findings with he localization of markers for different transcriptional activities which suggests a model where co-transcriptional splicing is accompanied by RPol stalling followed by super elongation complex driven resumption of transcription. The spatial arrangements of chromatin, nascent RNA, and transcription and splicing factors indicate a cooperative mechanism for co-transcriptional splicing hub biogenesis involving phase separation. Overall, our data and model suggest that splicing dynamics are regulated by local transcription factor concentrations.

## Results

### Imaging of co-transcriptional *kl-5* transcript splicing modalities and structures

To determine the spatial organization of transcription and splicing within the nucleus we performed RNA-FISH with an oligo-nucleotide probe that labels the microsatellite repeats of the giant introns of Y-loops A and C ((Fingerhut et al., 2019); Fig. 1A, B&C, Video 1), and with Hybridization Chain Reaction (HCR) probes for *kl-5* exon blocks 2-9 and 10-12. Both these exon blocks are less than 5 kb long. Exon 2-9 probes labelled a large region of the nucleoplasm substantially overlapping the labelling of the Y-loop A intron regions with the repeat probe. The exon 2-9 labelling shows a punctate, but contiguous ribbon pattern over several µm, which corresponds to hundreds of kB, much more than the extent of the genomic regions of these probes. This implies that exons are carried along the loop by the polymerase, which is inconsistent with a stationary transcription factory and similar to observations on transcription loops in a range of other metazoans including humans (Leidescher et al., 2022). Probes against the downstream exons 10-12 labelled a smaller portion of the loop and roughly overlapped in these regions with the exon 2-9 probes (the ellipse in the rightmost panel shows the region that was labelled by exon 2-9 probes, but not exon 10-12 probes). Thus, the part of the loop labelled only by the exon 2-9 RNA FISH probe corresponds to the genomic region of exons 2-9 and intron 9, and the portion co-labelled with the exon 2-9 and exon 10-12 probes is the loop downstream of exon 10. The two distinct, though overlapping, structures correspond to the two *kl-5* copies that these G2 phase cells contain (A and A’ in Fig. 1C). The intron probe labelling of Loop A (Fig. 1C, 1; Loop A is identified by the stronger AAGAC probe staining vs loop C which is not within the displayed volume, but can be seen in the accompanying video 1), overlapped with a large portion of the exons. However, a significant stretch of the loop that was stained by the exon probes was not labelled by the intron probe (ellipses in merged panel) indicating that all repeat containing introns were spliced out and the polymerase continued to transcribe while carrying only the exons. Thus, we show that splicing can be detected with this approach, and that the repeat probe-labelled *kl-5* introns 1, 9, 12 and 16 are spliced out co-transcriptionally, thereby supporting the findings of a study that showed co-transcriptional splicing of *kl-5* introns 1 and 16 (Fingerhut et al., 2024).

**Figure 1:**
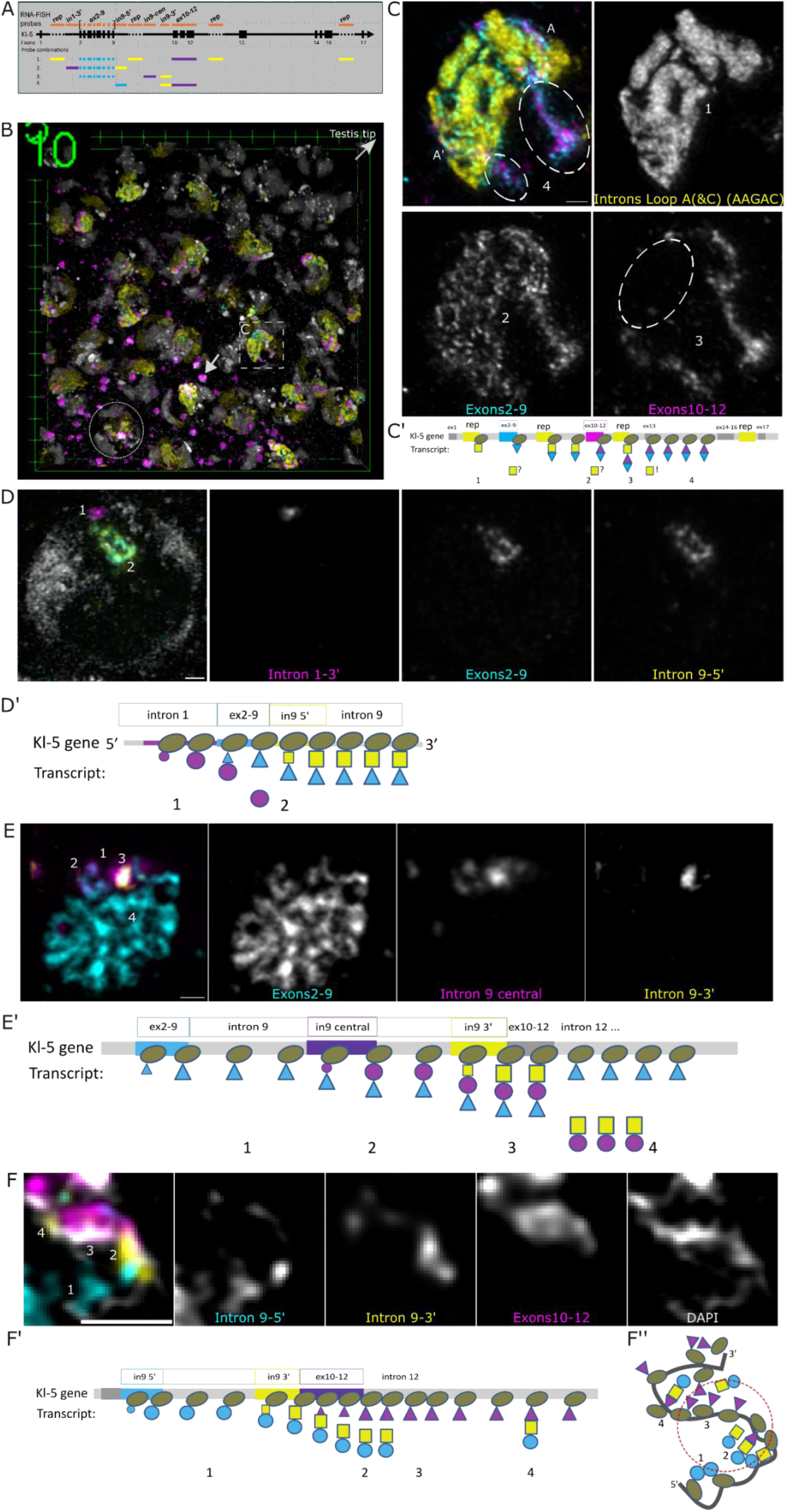
Co-transcriptional splicing. A) Structure of the *kl-5* gene, forming Y-loop A (black line; exons are indicated by black rectangles; dotted line indicates long repeat regions) and RNA-FISH probes applied in this study: FISH probe combination 1 consists of the AAGAC repeat oligo probe (rep) to label loop A introns strongly and loop C introns weakly, and HCR probe sets covering *kl-5* (loop A) exons 2-9 and 10-12. Combination 2: HCR probe sets for the 3’-region of intron 1, exons 2-9 and the 5’-region of intron 9. Combination 3: HCR probe sets for exons 2-9, the central region of intron 9 and the 3’-region of intron 9. Combination 4: intron 9 -5’, intron 9-3’ and exon block 10-12. B) 3D rendered overview of a z-stack of a whole-mount testis. Testis distal tip is towards the top right (arrow). Primary spermatocytes mature with increasing distance from the distal tip as reflected by increased Y-loop labelling and appearance of *kl-5* mRNA granules in the cytoplasm (magenta/cyan lacking intron signal; arrow). RNA-FISH staining corresponds to combination 1 in Fig. 1A; DAPI staining in grey shows the chromosome masses. One nucleus is indicated by a circle. Dashed rectangle C, see Figure 1C. Tick mark distance: 5 µm. Rendered volume: 77.16 x 77.16 x 14.28 µm (102 slices, vxl = 35 x 35 x 140 nm). C) Detail of Figure 1B (rectangle C) visualizes splicing of the *kl-5* transcript in a primary spermatocyte nucleus of a whole-mount testis. The ellipses in the merge (top left) panel mark Y-loop stretches labelled with exon probes 2-9 and 10-12 but not showing any repeat probe signal for the intervening intron, indicating that splicing has occurred. Loop C is not in the displayed volume. Scale bar: 1 µm. C’-F’) Schematics of our interpretation of transcription and splicing events along the *kl-5* gene. Indicated are the exons and introns which were probed. In all schematics ellipses indicate RPol, triangles indicate transcripts of exons, triangle corner down is exon block 2-9, triangle corner up is exon block 10-12. Squares and circles indicate introns. Colours correspond to the fluorescence images. In C’ the symbols right of the released (spliced) squares (introns) indicate whether we can conclude from these experiments if splicing has taken place, “?”: no conclusion can be drawn, “!”: splicing must have taken place by this point. 1: The repeat probe labels long intronic stretches throughout the *kl-5* gene (Y-loop A). 2: The exon 2-9 probe labels virtually the whole loop A, except for the first exon and intron, indicating that the RPol carries the transcript along the loop. 3: exons 10-12 only partially label the loop consistent with their transcription starting downstream of exon 9 and the long distance to exons 2-9, separated by a giant intron. 4: The repeat probe signal is lacking along a large portion of the loop that shows the signal of both exons, corresponding to RPols that carry both exon groups along intron 13 but had the repeat containing introns spliced out at exon 13, thus before the last exon and therefore co-transcriptionally. A small amount of repeat probe from intron 16 is detected in the large ellipse, but transcription has not yet reached this point in the smaller ellipse. D) RNA-FISH with probe combination 2. Co-labelling of an early loop in a stage 4 primary spermatocyte with exon 2-9 and intron 9 signals but absence of intron 1 signal reveals co-transcriptional splicing of intron 1. Greys in merged image (left panel) display DAPI. D’) 1: The 3’ end of intron 1 (magenta circle) is transcribed and accumulates locally, but not further downstream along the loop, due to co-transcriptional splicing. 2: exons 2-9 and the 5’ intron 9 probe co-label the rest of the early-stage loop. Scale bar: 1 µm. E) RNA-FISH with probe combination 3. The central intron 9 probe gives a more extended signal than the 3’ intron 9 probe, but both overlap, indicating that splicing occurs non-recursively. Video shows a z-scan through the whole nucleus including the FISH signals of the second *kl-5* copy. Scale bar: 1 µm. E’) 1: The exon 2-9 signal is transcribed and detected from there on. 2: This is followed by the central intron 9 signal. 3: The 3’ intron 9 region is transcribed and the whole intron 9 is spliced out shortly afterwards, thus leading to a local accumulation. 4: The exon 2-9 probe labels the whole loop since it is carried by Rpol to the end of the loop. F) RNA-FISH probe combination 4. Detail of the genomic region around exons 10-12, where intron 9-3’ and the following exons 10-12 are transcribed. Single optical section. Scale bar: 1µm. F’) 1: The 5’ intron 9 signal is detected as RPol transcribes intron 9. 2: The 3’ intron 9 signal is detected followed by the exon 10-12 transcript. The signal intensities are high at this stage, potentially due to an accumulation of spliced out introns that have not yet been processed. 3: The intron signals are lost indicating splicing has occurred. 4: Occasionally unspliced transcripts can be observed further downstream pointing towards delayed splicing. F’’) Schematic of transcription and splicing events along the chromatin fiber as seen in Figure 1F and corresponding Figure 1F’. Red circle indicates the main splicing region.

To determine transcription and splicing organization in more detail we improved the genomic resolution using HCR-RNA-FISH with probes for three short (<5 kb) consecutive regions to identify the exon 2-9 block and its flanking introns (Fig. 1D). Probes for exons 2-9 and the adjacent downstream 5’ region of intron 9 were overlapping over a large loop stretch as a chain of small blobs, confirming our previous observation that the polymerases carry the transcripts along the loop. Although we see separate blobs, we do not know the numbers of the RPols or transcripts per blob. Since the repeat intron probes used for the experiments shown in Fig. 1C cover large regions it was not clear whether RPol carries the introns or if they remained stationary relative to the template DNA. The contiguous Y-loop staining of HCR-RNA-FISH probes against the short 5’ intron 9 region showed that the intron RNA is carried by Rpol until it is spliced out (Fig. 1D). Initially focusing on S4 stage spermatocytes, we found the signal of a probe for the 3’ end of the preceding intron 1 (intron 1-3’) was highly localized with hardly any overlap with the other two probes. This confirmed that intron 1 was spliced out soon after its transcription and before transcription of the following intron, thus co-transcriptionally.

The extended length of the 3.2 Mb primary *kl-5* transcript would be about 1 mm, i.e. 50-100 times the diameter of the spermatocyte nucleus. Nevertheless, we see in Fig. 1C that the *kl-5* transcripts, labelled with the intron probe, occupy a discrete region within the nucleus and form a bulky loop structure suggesting they are confined close to the DNA loop (Fig. 1C).

Similarly, the exons closely follow the loop DNA fiber path. Obviously, processed exons, with the intervening introns spliced out, will be close to the transcribing polymerase but during intron transcription, due to the length of the introns, the 5’ exons could potentially be many µm distant from the polymerase. As we observe that the exon signal always follows a clear fiber path (e.g. Fig. 1D) and is close to the DNA fiber (e.g. Fig. 1F), it appears that either by folding of the nascent RNA, and/or by tethering, the 5’ exons are retained close to the polymerase. This may be important to bring the 5’ splice site into proximity with the 3’ splice site to facilitate efficient splicing.

### *kl-5* transcripts are spliced non-recursively

Carrying large transcripts likely generates a drag or accumulation of secondary nRNA structures that potentially interfere with the travelling RPol (Noe Gonzalez et al., 2021). The amount of carried transcript is strongly reduced by co-transcriptional splicing, but each of the giant introns is still up to several 100 kb long, thus suggesting that splicing might take place already while they are being transcribed, i.e. as recursive splicing (Shepard et al., 2009). To detect recursive splicing, we performed HCR-RNA-FISH with 2 probes for intron 9, the central and the 3’ regions, together with probes for the preceding exon 2-9 block (Fig. 1E).

Recursive splicing would be indicated by a separation of the intron probe signals. The central intron 9 probe labelled an extended stretch of the exon 2-9 labelled loop and accumulated locally, overlapping with the 3’ intron 9 probe, indicating that the central region of intron 9 is retained in the nRNA when the 3’ region is transcribed. This provides evidence that intron 9 is completely transcribed before being spliced out, thus non-recursively. To unambiguously identify where splicing of intron 9 occurs, the exon block following intron 9, comprising exons 10-12, was labelled together with 5’- and 3’-probes of intron 9 (Fig. 1F).

From the DAPI labelling of the chromosome fiber and proximal vs. distal loop FISH labels the direction of transcription can be inferred. Proximally the fiber only carried the upstream intron region label (intron 9-5’), then more distally there was an accumulation of the 3’ intron probe (intron 9-3’), at which point the downstream exon probe label was detected (exons 10-12). From that point on both intron labels strongly decreased, thus confirming co-transcriptional non-recursive splicing of the *kl-5* intron 9. The splicing activity localizes to a region of low-density chromatin coming off the main fiber, towards the concave side within a fiber bend. This potentially identifies a splicing ‘hub’ which is further investigated below.

### Splicing decisions are made on a cluster-by-cluster basis and are frequently delayed in more mature spermatocytes

When analyzing the distribution of RNA-FISH signals along Y-loop A in more mature spermatocytes (S5/6 stage) we frequently found signals of the intron 1 3’ end covering a large part of the loop, and overlapping labelling of the downstream exon (exons 2-9) or intron probes (intron 9-5’). Labelling of both intron probes was lost further downstream along the loop, while the exon probes were retained, thus indicating delayed co-transcriptional splicing (Fig. 2A). We estimated the splicing delay of intron 1 by measuring the distance along a loop covered by FISH signal. Most values were between about 2 and 6 µm, and up to 10 µm, corresponding to hundreds of kB (Fig. 2). The example in Figure 2A shows a late-stage spermatocyte where intron 1 was detected along a 6 µm fiber stretch (Fig. 2 A). Even though intron 1 and 9 signals overlapped for such long distances, they were not spliced out together, as intron 9 was still found further down the loop (Fig. 2A).

**Figure 2:**
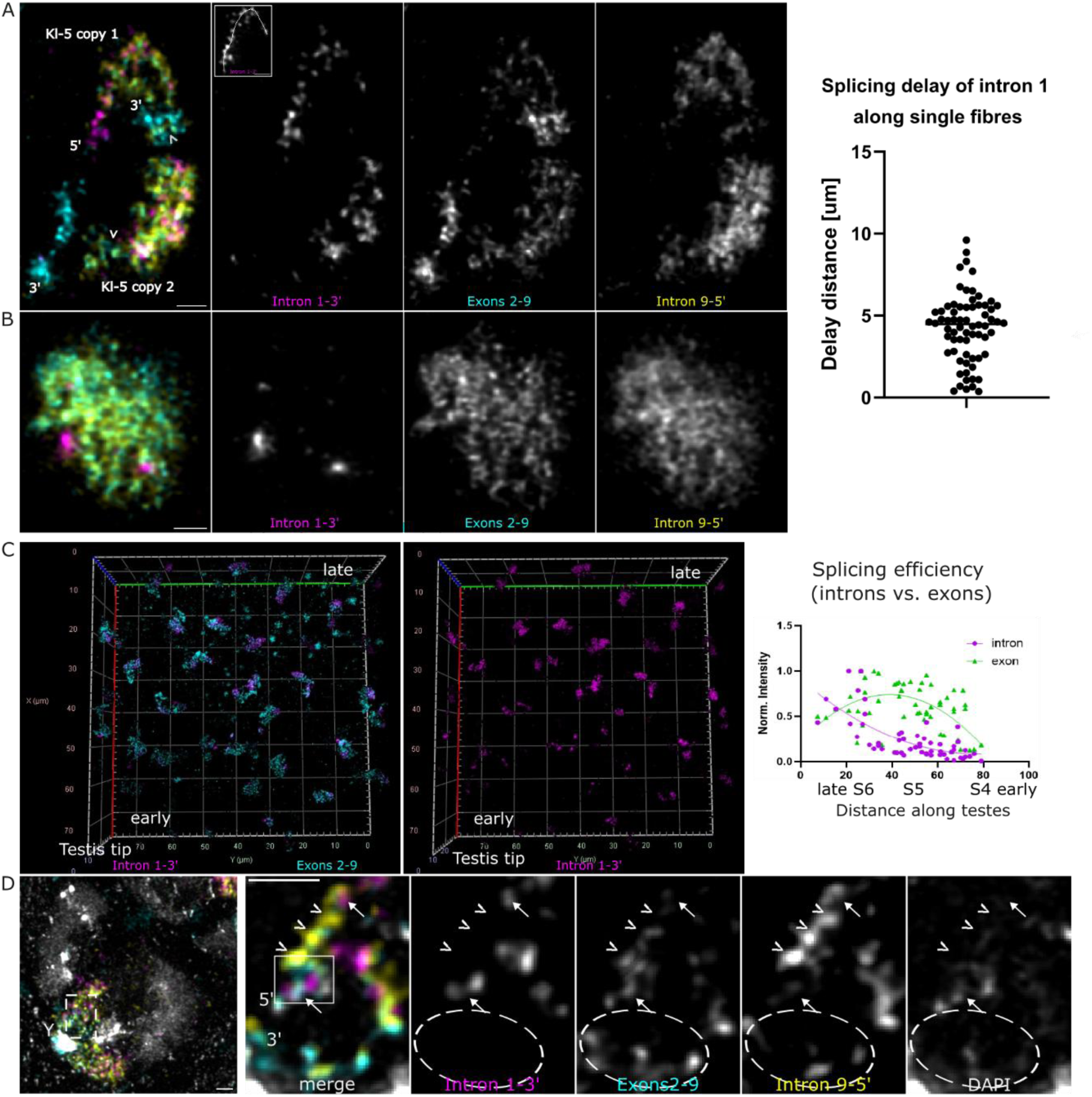
Delayed intron 1 splicing increases with spermatocyte maturity. RNA-FISH probe combination 2 in all panels. A) The intron 1-3’ probe is retained for considerable lengths along Y-loops of late stage primary spermatocytes (S5-6). Intron 9 is not spliced out concomitantly with intron 1 (intron 9 with exons 2-9, without intron 1; arrowheads). Eventually both introns are spliced out as indicated by the Y-loop fibers solely stained with the exon probe (towards the 3’ ends). Intron 1 and exons 2-9 without intron 9 label the 5’ proximal region. The 5’ proximal region of *kl-5* copy 2 could not be unambiguously identified. Maximum intensity z projection of 28 optical slices (140 nm / slice, total 3780 nm). The insert indicates the line drawn for splicing delay estimates. Fiber lengths labelled with the intron 1-5’ probe are measured. In this example: 6.2 µm. Scale bar: 1 µm. “3’” at the *kl-5* copy 1 of the merged image (left panel) indicates directionality of the loop and is intended for orientation only – the actual 3’ end is close to the 5’ end, where the intron 9-5’ label is found. Scale bar: 1 µm. B) In contrast to the later spermatocytes shown in A), in stage 4 primary spermatocytes intron 1-3’ accumulates in one intense spot per gene copy indicating rapid splicing. The exon 2-9 and intron 9-5’ probes are spread throughout the nucleoplasm. Maximum intensity z projection of 25 optical slices. Scale bar: 1 µm. Graph shows the distributions of intron 1 delay distances along Y-loop fibers as indicated in the inlay of Fig. 2A. 96 measurements of a variety of spermatocyte stages in 2 testes. C) 3D rendering of a whole-mount testis volume containing early and late-stage primary spermatocytes. Later spermatocytes (towards top right) contain more intron 1 signal, absolute and relative to exon 2-9 signal. Graph shows the total intensity of intron 1 (magenta circles) and exon 2-9 (green triangles) signals for each nucleus as a function of its distance from the most mature spermatocytes. The nuclei furthest away from the point of highest maturity (earlier stages, S3/4) are low in Y-loop transcript. Exon intensities (green) sharply increase as they mature (towards the left in the graph) until they reach a plateau at the S5 stage and then decrease, due to ceased transcription initiation and export of processed mRNA from the nucleus (as seen by cytoplasmic exon particles) at the S6 (most mature primary spermatocyte) stage. D) Crop of a nucleus with extensively delayed intron 1 splicing. Left: merged overview of the whole nucleus (maximum intensity projection of 38 slices, dz = 140 nm, 5.18 µm thickness). The Y-chromosome is indicated by “Y” and the directionality of the *kl-5* loop by “5’” and “3’”. In the detail (rectangle in the overview; maximum projection of 4 slices) two regions along the loop can be distinguished according to their coverage or depletion of intron 1-3’ signal respectively (ellipse). Stretches of unspliced clusters (arrows) are interspersed by clusters where intron 1 was spliced out (arrowheads). The rectangle indicates an area of intron 1 splicing. Scale bar: 1 µm.

Interestingly, this delayed splicing was not a constant feature of all nuclei but varied with the developmental stage of the spermatocytes; it was rare in early (stage 4; Fig. 2B), but frequent in late (S5-6; Fig. 2A) spermatocytes. To correlate splicing delay with spermatocyte development we measured probe fluorescence intensities of intron 1-3’ and exon 2-9 HCR-RNA-FISH probes in nuclei along the proximal-distal testis axis, which reflects spermatocyte developmental time. The intron probe target region is right before (5’ of) the exon probe sequence and of similar size (3.4 and 2.2 KB), so the lag of transcription would be in the range of minutes, while the scale of the graph is hours to days. If co-transcriptional splicing was efficient, intron intensity would remain low. Conversely, the intron signal would be expected to increase at the same rate as the exon signal if it was consistently retained.

However, in contrast to the steep increase of exon 2-9 probe intensity in S4 cells, intron 1-3’ probe intensities remained initially low from S4 to S5 and only then started to increase (from right to left in the graph). This shows that co-transcriptional splicing is efficient in S4 cells and delayed in more mature primary spermatocytes.

To obtain insight into the regulation of delayed splicing we asked if splicing is localized to a specific position along the Y-loop, or whether it is related to a more dynamic, stochastic process, e.g. concentration dependent (suggested by e.g. (Ullrich et al., 2025)). For this, we determined the relative positions of intron 1 spliced and un-spliced clusters along the same chromatin fibers of late-stage spermatocytes with extensive delayed splicing and found stretches of unspliced clusters (containing intron 1-3’) interspersed with spliced clusters (only containing exons 2-9 and intron 9-5’) (Fig. 2D). Taken together, this shows that splicing decisions are made on a cluster-by-cluster basis and is delayed with increasing spermatocyte maturity. As also noted in Fig. 1F and Fig. 3, a bend indicative of a splicing hub is formed around splicing events (as identified by the transition from unspliced to spliced clusters; rectangle in Fig. 2D).

**Figure 3:**
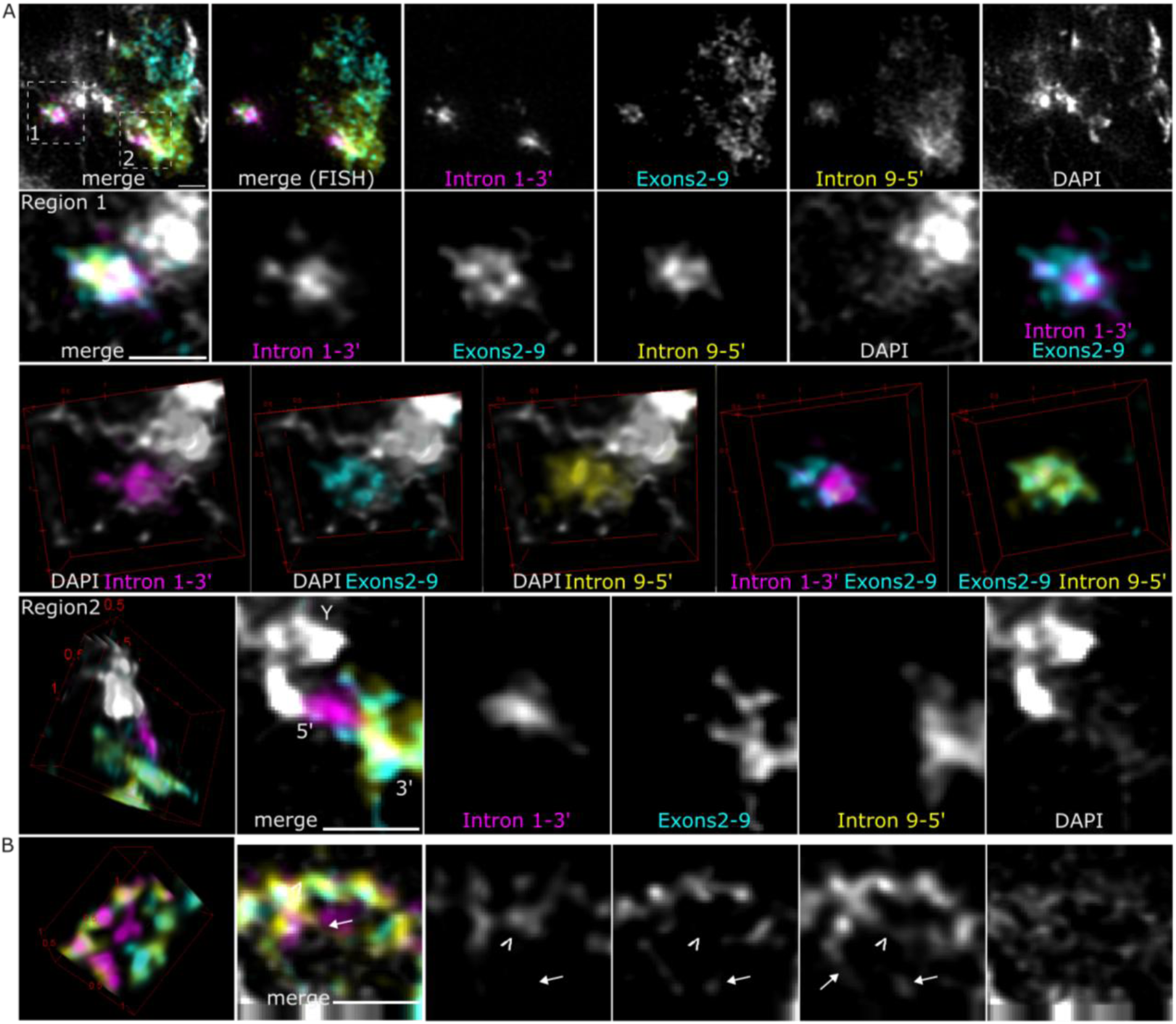
Hubs. A) shows a nucleus of an early to mid-stage spermatocyte with little delayed intron 1 splicing. RNA-FISH probe combination 2: intron 1 -3’, exons 2-9, intron 9-5’. Top row shows a cropped nucleus region (average intensity projection of 9 slices, 3.2 µm thickness). The two strong intron 1 signals indicate the 5’ loop A regions of the two *kl-5* copies. The transcriptional progress of the two copies differs considerably, as judged from the amounts of exon block 2-9 and intron 9-5’ signal. Rows 2 and 3 show details of splicing region 1 (rectangle 1 in top row) where spliced transcripts have the exon 2-9 signal in the absence of intron1-3’. Exons 2-9 (and thus presumably the path of the RPol) forms a horseshoe shape as visualized by 3D rendering in row 3 panel 2. Intron 1 accumulates centrally at the splice region, wrapped by DNA fibers. The 5’ intron9 label is enriched between them. Row 2: Average z projection of 9 slices (same volume as in A). Row 3: 3D rendering of the first three panels with DAPI, then intron 1-5’ and exon 2-9 probe signals, and exon 2-9 and intron 9-5’ signals. Distances are indicated on the box (500 nm/tick). Row 4 shows region 2, the 5’ region of the further transcribed loop A copy, whose structure is more extended. B) Intron 1 signal can be found enriched off the main fiber axis (arrowhead in panels 3-5), on small chromatin loops between nearby fibers along which exon 2-9 and intron 9 signal, partially together with intron 1 signal, is arranged. A bit further along the fiber are exon 2-9 and intron 9 signals without intron 1 (arrow), indicating that intron 1 was spliced out, likely at the accumulation between the fibers, where it is less closely associated with the other FISH signals (most obvious in the 3D rendering of FISH signals (left panel)). Other panels: maximum intensity projection of 2 slices. Scale bar: 1 µm.

### Co-transcriptional splicing takes place in hubs

Fig. 3 illustrates architectures associated with splicing. The splicing events can be localized to the region where the exon 2-9 signal loses the overlap with the intron1-3’ signal. The DAPI label here shows decondensed DNA emanating from the highly condensed heterochromatic Y-chromosome mass, and the exon 2-9 and intron 9-5’ signals show that splicing is associated with a tight fiber bend around a blob of intron 1-3’ signal, potentially forming a ‘hub’ with local concentration of splicing events (Fig. 3A, region 1). This likely indicates the beginning of the loop and suggests that loop formation is driven by transcription (consistent with work from others, including (Caudron-Herger et al., 2011; Leidescher et al., 2022).

The exon 2-9 probe appears as small blobs along the DNA, while the intron 9-5’ probe is enriched between the exon and intron 1 probes, even though intron 9 has been transcribed more recently, indicating a strong association of exon-nRNA with the polymerase, in line with a mechanism that ensures proximity of consecutive exons, as shown by cryo EM structures for RPol and the spliceosome component U1snRNP (Zhang et al., 2021).

We also note, when comparing the transcripts of the two *kl-5* copies within the same nucleus we found a strong difference in transcriptional progress (Fig. 3A, regions 1 and 2). The intron 1-3’ accumulation of the more progressed *kl-5* loop is more elongated (region 2), highlighting varying structure and dynamics at corresponding genomic locations of the two *kl-5* copies.

Fig 3B shows another variation of a splicing hub in a more mature nucleus. Here, intron 1 signal is enriched in the center of intertwined Y-loop A fibers, only loosely associated with exon signal.

Overall, we show that co-transcriptional splicing is organized in hubs of varying arrangements with spliced introns accumulating between fibers.

### dEaf localizes in primary spermatocytes in nuclear speckles that exhibit phase separation properties

Splicing and transcription are mechanistically and dynamically linked (Bentley, 1999; Caizzi et al., 2021; Fingerhut et al., 2024). We have previously shown active and inactive chromatin marks indicating phases of varying transcriptional activities along Y-loops (Koestler et al., 2024). Cdk9 activity is required for transcription pause release by phosphorylation of PSer2 of the RPol CTD, and for efficient splicing (Egloff, 2021). Since Cdk9 is often incorporated into the Super Elongation Complex (SEC) (Luo et al., 2012b), we made use of a fly line from the CPTI collection (CPTI-000560, (Lowe et al., 2014)) expressing endogenously YFP-tagged Eaf to further investigate transcription dynamics along the Y-loops (Lye et al., 2014) (Fig. 4A). Eaf is a SEC component for which nuclear expression in *Drosophila* embryos has previously been observed. Normal functioning of the tagged protein was indicated by the observation that YFP-Eaf flies produced healthy offspring, while Eaf is essential for development (Lye et al., 2014; Smith et al., 2008). Confocal microscopy imaging of whole-mount testis volumes comprising germ cells at the testis tip, up to late-stage 6 primary spermatocytes showed a lack of Eaf in early cells. Diffuse signal started to appear in spermatogonia (Fig. 4A). In S3 spermatocytes Eaf speckles began to accumulate at the nucleolar periphery. At the S4 stage Eaf largely encapsulated the nucleolus, accumulated in chromosome masses and in speckles in the nucleoplasm. This pattern persisted, except that in late S5/S6 stages the overall Eaf intensity decreased. Eaf was absent from somatic cells in the testes.

**Figure 4:**
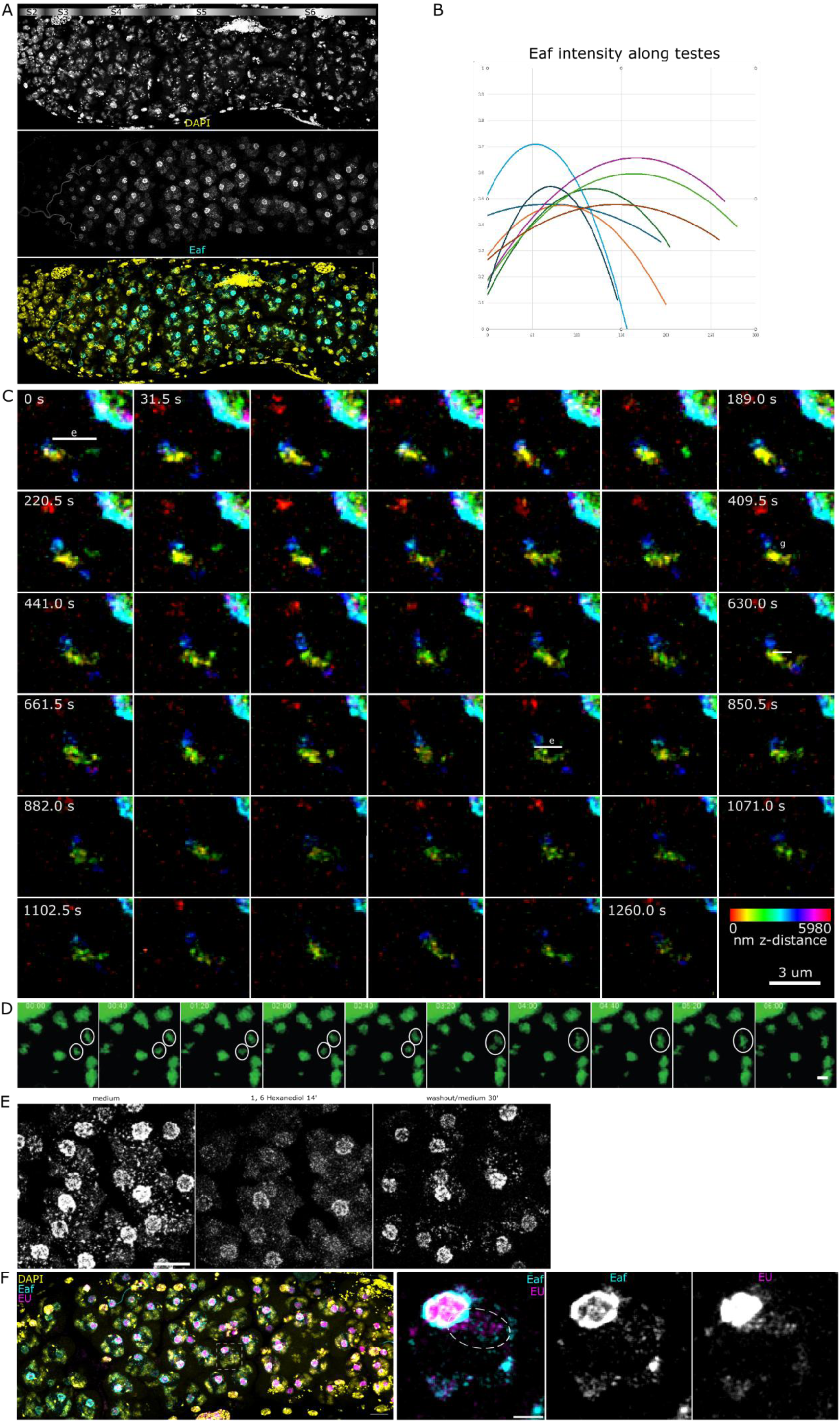
Eaf in *Drosophila* testes. A) Distribution of Eaf in *Drosophila* testes. Yellow, and top panel: DAPI; Cyan and middle panel Eaf. Distal testis tip is to the left. The bar on the top provides a guide for the progression of primary spermatocyte stages across the testis. Maximum intensity projection of z=146 slices, dz=346 nm, covering 50.2 µm, approximately half the testis volume. Scale bar: 10 µm. B) Graph shows the Eaf intensity (y-axis) along testes (from nearby the distal tip towards further proximal; x-axis). Shown are trend-lines of 8 line plot measurements in 6 testes. Raw data with the measured regions indicated are provided in the Supplementary Figure. C-D) The super elongation complex component Eaf is dynamic and forms speckles, consistent with phase separation properties. Live imaging of YFP::Eaf in primary spermatocytes of whole-mount testes. C) shows the re-arrangement of an Eaf speckle over 21 min with a frame rate of 31.5 s. A timelapse sequence of a depth-color-coded maximum intensity projection of a 6 µm z-stack of a ROI within a larval testis is shown. The nucleolus is at the top right corner. The shape of the central Eaf speckle shifts between globular and elongated/filamentous, suggesting a correlation between Eaf speckle structure and transcriptional activity, e.g. bursts and pauses. The association of Eaf with chromatin (Fig. 5) suggests its involvement in the observed re-arrangements. Scale bar: 3 µm. D) Eaf speckles move and fuse. Circles indicate 2 Eaf speckles that move and eventually fuse. Timelapse, 40s frame rate. The nucleolus is in the top left corner. Scale bar: 1 µm E) Eaf speckles are sensitive to 1, 6 Hexanediol and reform upon wash out. Maximum projection of a z-stack of 33 slices with a distance of 1 µm. A time-lapse movie was acquired with a frame rate of 2 min. The 3 panels show primary spermatocytes in a whole-mount larval testis expressing Eaf-YFP. Left: in medium, before addition of 1, 6 Hexanediol. Centre: after 14 min in 5 % Hexanediol, and right, 30 min after washout. Scale bar: 10 µm. F) Nascent RNA accumulates adjacent to Eaf speckles. Overview: Maximum intensity projection of 46 slices (dz=500 nm). Merged image of DAPI (yellow), Eaf (cyan), and EU labeling (magenta). Testis tip is to the right. Scale bar: 10 µm. Detail panels show the boxed nucleus in the overview. Merge of Eaf (cyan) and EU (magenta) and single channels in grey. Indicated (ellipse) is a region where Eaf and nRNA (EU) accumulate, most likely at a Y-loop emanating from the Y-chromosome which is located next to the bright nucleolus. Maximum intensity projection of 5 slices. Scale bar: 3 µm.

YFP-Eaf frequently appeared as nuclear speckles with diameters up to 1-2 µm but also much smaller, consistent with the characterization of human EAF1 as a component of Cajal bodies (CB; Polak et al., 2003), phase separated nuclear structures containing spliceosome components (Fig. 4C, D, E, F and Videos 2-4). The SEC can phase separate (Guo et al., 2020), and EAF1 enhanced phase separation of ELL is necessary for transcriptional restart after DNA damage repair (Pal et al., 2026).

Live imaging of YFP-Eaf in whole-mount testes showed Eaf speckle movement within the nucleoplasm and fusion events (Fig. 4C, D and Videos 2, 3 and 4). Most Eaf-YFP speckles within a 30 min observation time frame wobbled around a rather fixed position, with a few events showing movement quite directly towards the periphery. Events of splitting and fusion were also observed (Fig. 4D, Video 4).

Treatment with 5 % 1,6-Hexanediol dissolved Eaf speckles with effects being observable after less than 5 min and full dispersal in most nuclei within 10 min (Fig. 4E). Speckles started to reappear within about 5 min after washout and largely returned after 10-20 min. Recovery within the observed time of 30 min was not complete, also partially due to photobleaching of YFP (Fig. 4E and Video 5). Although the effect of 1,6-Hexanediol could be indirect through its effect on kinases these results are consistent with liquid-liquid phase separation and suggest that *Drosophila* Eaf behaves similarly to its human homolog (Alberti et al., 2019; Düster et al., 2021; Pal et al., 2026).

EU-Click labelling showed nascent transcription adjacent to Eaf, and further extending from there, suggesting Eaf-, and thus presumably SEC-, dependent Cdk9 activity at the periphery of speckles (Fig. 4F). This is consistent with the localization of ELL and EAF1 in CBs suggesting that targets for the transcriptional elongation activity of ELL might exist either within or adjacent to CBs (Polak et al., 2003), and with a model for RPol pause release at the periphery of phase condensates (Pancholi et al., 2021).

### The spatial arrangement of RPol, spliceosome, Eaf-speckles and chromatin suggests a cycle of transcription, pause/splicing, and pause release at splicing hubs that induce chromatin bends

To better understand co-transcriptional structure/dynamics relationships and regulation we performed high-resolution immunofluorescence studies using Zeiss Airyscan 2 microscopy with joint-deconvolution (Prazeres da Costa et al., 2021). It showed association of YFP-Eaf with splicing factors (anti-Sm, targeting the Sm proteins that form the core of the spliceosome’s snRNPs) and transcribing RPol, RPolPSer2 (Fig. 5A), thus further corroborating the involvement of the SEC with co-regulation of splicing and transcription, and consistent with the finding that the hyperphosphorylated C-terminal domain of RPol is preferentially incorporated into condensates that are formed by splicing factors (Guo et al., 2019). This suggests that the SEC accumulates at transcription elongation pause/splicing regions to phosphorylate Ser2 of paused RPol which then proceeds with transcription, leaving the pause site (consistent with (Pancholi et al., 2021)).

**Figure 5:**
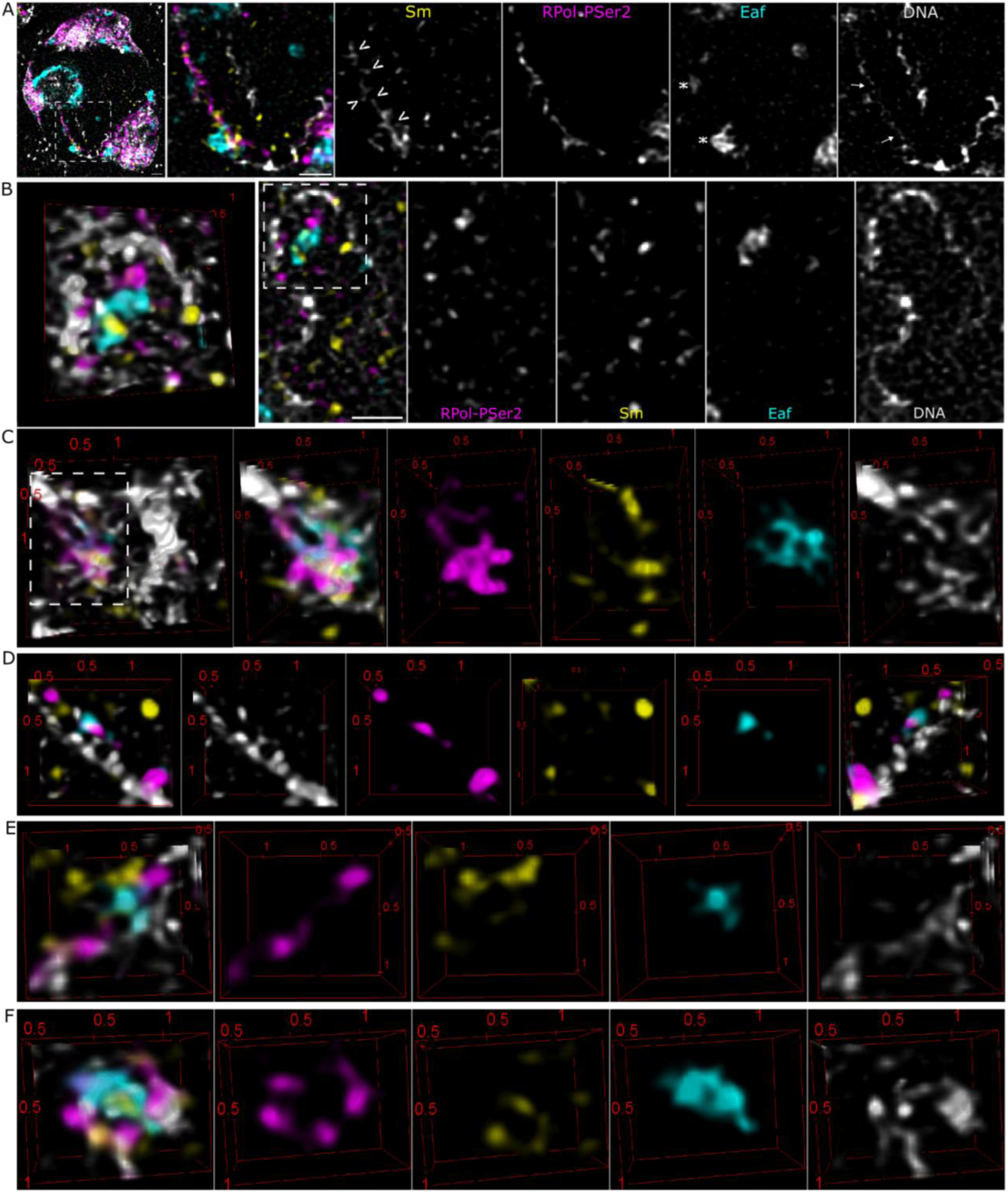
The spliceosome, elongating RNApol and the super elongation complex component Eaf are arranged along Y-loops. Primary spermatocyte in a whole-mount testis, immunostained against the Smith antigen labelling the major spliceosome proteins (Sm; yellow in merged images), Phospho-serine 2 of the RNA polymerase II CTD repeat indicating active transcription (magenta); Eaf was visualized by the fluorescence of the YFP tag (cyan), and DNA by DAPI (grey). A) Shows a transcriptionally highly active chromatin fiber (arrows in DNA panel), occupied by elongating RPol over several µm, and associated with spliceosome components (Sm; arrowheads). Eaf is associated with the fiber but more localised and off the main axis of the fiber (asterisks) Single optical section. Scale bar: 1 µm. B&C) show enrichment of RPol/Sm/Eaf on decondensed chromatin fibers, off a dense, bent chromatin fiber, on the concave side, forming a splicing/transcription hub. B) left panel shows a 3D rendering of the boxed region in the adjacent merged image. C) 3D renderings of another hub example adjacent to a bent condensed chromatin fiber, included in the crop in the left panel. D)-F) Structural arrangements suggest different stages of the process of splicing hub formation. D) A straight fiber from which short fibers (loops) protrude, with small local enrichments of elongating RPol, often together with Eaf or Sm. E) A moderately bent fiber along which elongating RPol is localized as well as Eaf and Sm (both off the main chromatin fiber axis). F) Two fibers (potentially a bend) with RPol arranged around a bigger Eaf blob. Sm is in the center of the Eaf blob without RPol, and with elongating RPol along the fiber, potentially reflecting two different functional spliceosome states. Distances are indicated on the box frame in µm.

More specifically, RPol was generally localized along decondensed chromatin, and often together with spliceosome components (Sm) as indicated in Figure 5A and B. Whether this corresponds to co-transcriptional splicing or only the spliceosome component U1 snRNP, which can directly interact with RPol and the 5’ splice site of nRNA (Zhang et al., 2021) remains to be shown. Eaf speckles ranging from about 0.1-1 µm in diameter were associated with both RPolPSer2 and Sm. Large accumulations were arranged in hubs on the concave side of fiber bends (Fig. 5 B&C). Overall, the relative arrangements of Sm, RPolPSer2 and Eaf along Y-loops as well as the associated chromatin structure varied, likely reflecting different phases of the transcription elongation – pause/splicing – pause release cycle. Fig. 5 D-F show the putative three main steps/arrangements: step 1 (Fig. 5D): transcription: along mostly straight chromatin fibers single separate spots of RPol-PSer2 were located on small bulges off the main fiber axis, presumably representing small loops as we previously showed by super-resolution STORM and STED microscopy of isolated spermatocytes (Ball et al., 2023).This suggests a lampbrush like organization, with small decondensed chromatin loops protruding from the main axis to allow transcription and implicates a highly dynamic process of loop extrusion and back reeling while RPol transcribes. Also, single spots of RPolPSer2 were often associated with small spots of Sm signal and small Eaf speckles. Step 2, initiation of hub formation (Fig. 5E): Several RPolPSer2 clusters, Sm and Eaf speckles form a loose aggregate along a bent chromatin fiber. Step 3, an established hub (Fig. 5F): Large accumulations of RPolPSer2 and Sm associated Eaf speckles on the concave side of strongly bent fibers, like the splicing events that we observed by RNA-FISH (Fig. 1F, 2D and 3A&B) with RPolPSer2 along the chromatin fiber and the Eaf speckle bridging the fibers. Sm is either associated with RPolPSer2 or more central within the Eaf speckle, suggesting different states, the first before or during splicing, the latter after splicing, before spliceosome/Rpol assembly.

Imaging of paused RPolPSer5 also showed its association with spliceosome components and Eaf (Fig. 6). Detailed analysis of the spatial relationship of Eaf with RPolPSer2 and RPolPSer5 by segmentation of Eaf and RPol volumes showed that Eaf was more closely associated with RPolPSer5 than with RPolPSer2: the relative fraction of Eaf overlapping with RPolPSer5 was over 10 % vs around 4 % of RPolPSer2 (Fig. 6 box plot). This is consistent with the experiments and model with RPolPSer5 inside the phase condensate and RPolPSer2 at the periphery from Pancholi et al. (Pancholi et al., 2021).

**Figure 6:**
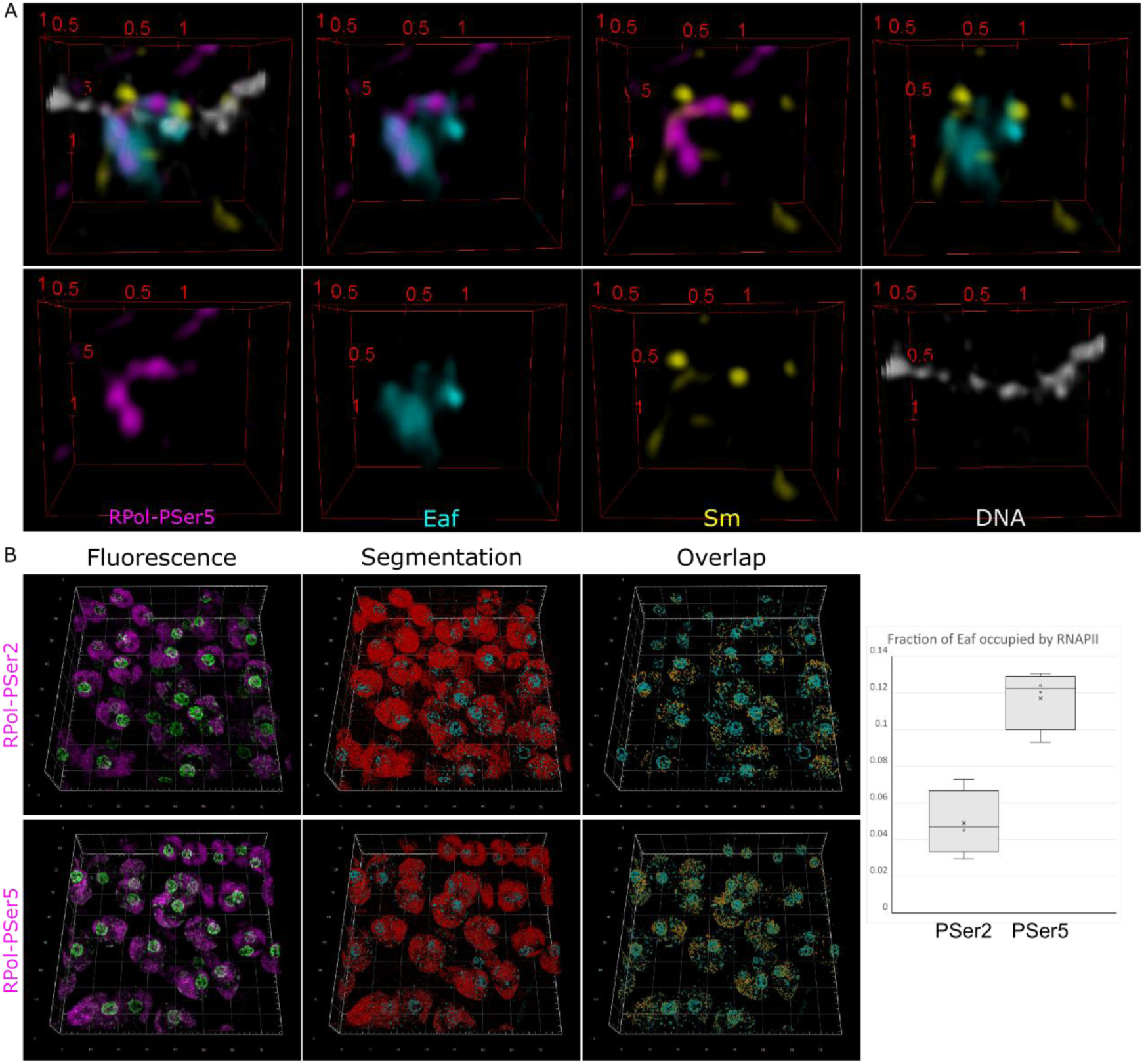
Engaged/pausing RNA polymerase II (RPolPSer5) is more strongly associated with Eaf than elongating RPolPSer2. A) shows an example of RPolPSer5 (magenta) along a chromatin fiber (DAPI, grey), together with Eaf (cyan) and Sm (yellow). 3D renderings. Distances are indicated on the box frame in µm. B) Quantifications of Eaf associated with pausing (RPolPSer5) and elongating (RPolPSer2) RPol. Shown are 3D renderings of fluorescence images, their segmented volumes and the Eaf/RPol overlap. Plot shows the volume fractions of Eaf that are occupied by RNAPII-PSer2 or -PSer5, respectively. 4 independent samples (testes) per staining. T-test (2-tailed, heteroscedastic) gives a p-value of 0.001. Box plots indicate 1^st^ and 3^rd^ quartiles, the median, and local minima and maxima.

These results suggest a transcription pause initiated phase separation mechanism of concentrating nascent RNA, RNA binding proteins and transcription factors for efficient splicing and pause release, and shaping chromatin fiber structure by inducing curvature.

## Discussion

In this paper we make two major observations that are interrelated. We show that co-transcriptional splicing delay increases with spermatocyte maturity, and we provide a model for splicing hub biogenesis, based on the presented data and a wealth of previous studies.

Variations in splicing kinetics have been shown in numerous instances. A majority of genes are spliced co-transcriptionally, but intron retention is also an important mechanism of protein abundance control, especially during development, but also in response to external factors (Porras-Tobias et al., 2025). There is evidence for inversely correlated coupling of transcription rate and co-transcriptional splicing efficiency (Khodor et al., 2012) and it has been suggested that highly expressed genes show more splicing delay, potentially due to competition of splicing factors (Ullrich et al., 2025). Also, different introns within the same gene can be spliced with different efficiencies indicating local factors in gene architecture to be important, and even the same intron can be delayed to varying extents pointing to external factors (Khodor et al., 2012).

Our observation of a systematic increase in splicing delay with spermatocyte maturity provides an entry point to determine the factors involved. Here we provide evidence for the involvement of Eaf in the context of a model linking splicing to transcriptional pause/release. Previously, it has been shown that transcriptional elongation and pause can take place in different types of condensates (Guo et al., 2019), that transcriptional pause can be associated with splicing (Mayer et al., 2017; Shine et al., 2024), and that spliceosomes are assembled in phase separated organelles (Cajal bodies; (Hyman et al., 2014)). Here, by using Airyscan microscopy enhanced with joint-deconvolution we show how elongating and pausing RPol, spliceosomes and the super elongation complex (Eaf) are arranged relative to each other and to chromatin. Our evidence for transcriptional pausing along Y-loops consists of the localization of the SEC factor Eaf and RPolPSer5. Eaf indicates a requirement for RPolSer2 phosphorylation, thus implying the occurrence of the RPolSer2 unphosphorylated non-elongating form. Similarly, RPolPSer5P is commonly associated with promoter proximal pausing. From the different spatial arrangements of splicing and transcription factors, we were able to deduce a dynamic model of splicing hub biogenesis through intermolecular forces between chromatin, RPols, nRNA, splicing and transcription factors (Fig. 7, box). In this model transcriptional pause of an RPol cluster either due to splicing, a checkpoint or condensed chromatin along a chromatin fiber leads to an accumulation of RPols. Nascent RNA and RNA binding proteins recruit spliceosomes, and after splicing SEC activity releases RPols from the pause. The interaction of involved proteins and nucleic acids draws them towards a common centre, shaping chromatin and forming a hub, which increases the local concentrations of splicing and transcription factors, potentially via phase separation. This is consistent with previous findings which show that splicing factors copurify preferentially with RPolPSer5 which can form phase-separated condensates where splicing factors are concentrated (Shenasa and Bentley, 2023), and our imaging experiments presented here showing the spatial arrangements of transcription and splicing factors and chromatin along with the live imaging of 1,6-Hexanediol treated testes. We also show that transcription takes place off the main, more condensed chromatin fiber, most likely on small loops, in a lampbrush-like manner. This implies constant extrusion and retraction of these loops in waves as the RPols travel along the fiber. The off-axis arrangement also allows interacting transcription clusters to induce fiber bending during hub formation. Upon splicing completion pause release allows RPols resumption of elongation and either hub dissolution or retaining a more stable hub core that provides components for oncoming RPol clusters. Our model suggests a feedback loop of accumulating splicing factors, splicing product and approaching RPols, that allows quick and transient hub formation, thus enabling flexible tuning of transcription elongation vs. pause/splicing kinetics.

**Figure 7:**
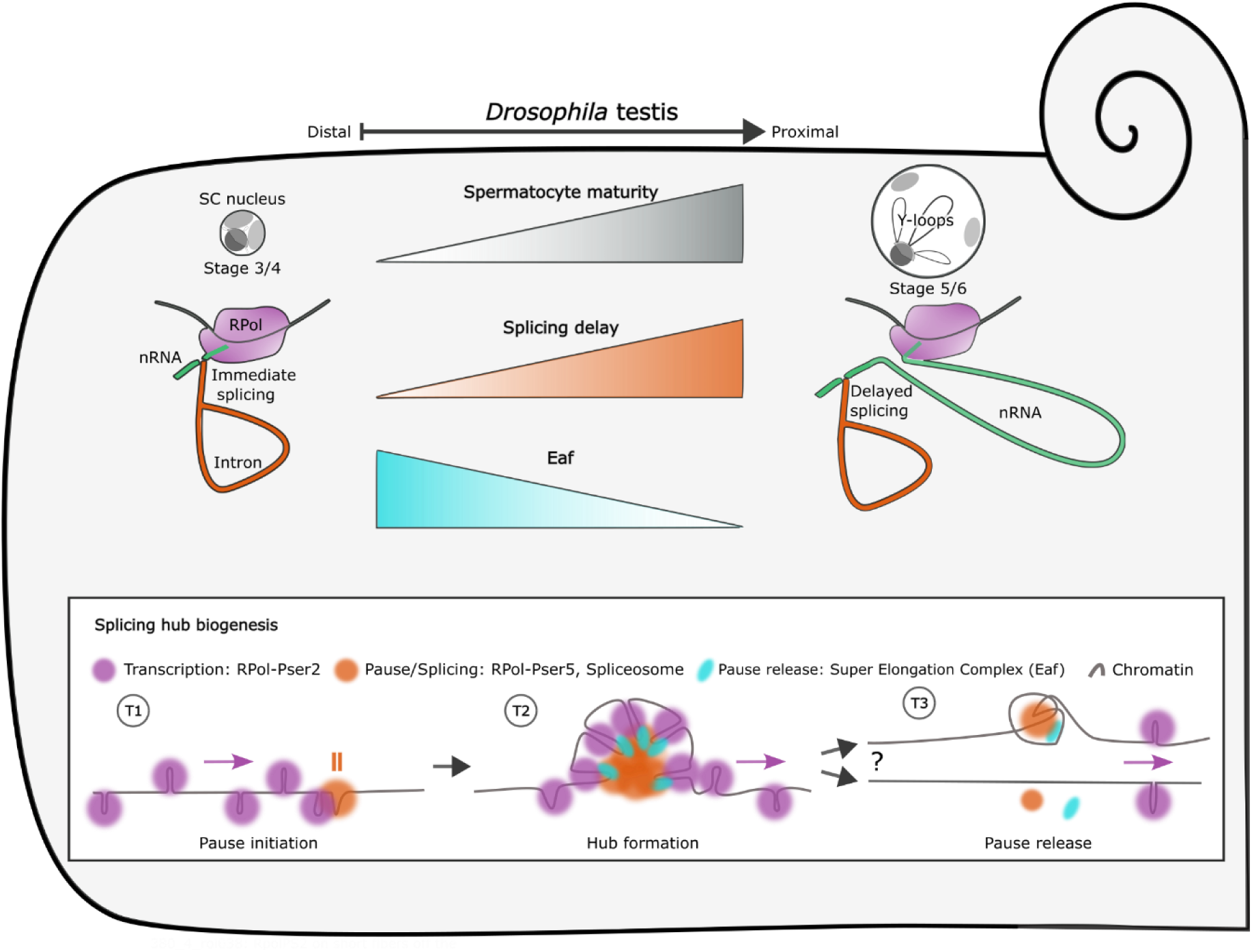
Summary diagram. In *Drosophila* testes, spermatogenesis is initiated distally near the tip and progresses towards their proximal side. Splicing delay increases with spermatocyte maturity and is inversely correlated with Eaf concentration. SC: spermatocyte. Box - Splicing hub biogenesis: Timepoint T1: RPol clusters travel along the chromatin fiber while carrying and transcribing nRNA from continuously extruding and re-folding loops off the main, more condensed, chromatin fiber axis until they reach a pause point. The nature of the pause signal is not clear: the stochastic distribution of delayed splicing indicates a multi-parametric process, potentially resulting from the amount of produced nRNA and local concentration of splicing factors, or chromatin structure and nucleotide composition. RPol-Ser5 is phosphorylated and spliceosomes assembled. T2: Pausing allows several RPols/Splicesomes to form a hub with a core of paused RPol-PSer5, spliceosomes and spliced intron RNA causing the chromatin fiber to bend. The super elongation complex (observed by its component Eaf in the experiments) releases the RPol from pause after splicing and a potential checkpoint control. T3: Once a train of RPols has left the hub, splicing and transcription factors and the affected chromatin structure may be retained to enable quick reformation during arrival of the next RPol train, or dissolved.

The decrease in Eaf concentration that we observe during spermatocyte maturation, which is probably at least partially due to the increase in nuclear volume of about 25 times, therefore provides a link to the corresponding increase in splicing delay. It suggests a decrease in splicing hubs and therefore more time until an RPol reaches one and vice-versa the decrease in RPol density leads to fewer hubs. A decrease in pausing for splicing might be a mechanism to increase local transcription rates of the trailing RPols as meiosis approaches. This is consistent with the observation that splicing can become rate-limiting when transcription is fast (Carrillo Oesterreich et al., 2016).

The stochastic distribution of splicing hubs that we observe along the same fiber also suggests a concentration dependency rather than fixed positions e.g. by specific sequences as suggested by (Liao and Regev, 2021) and indicates that they are formed temporarily and then again dissolved. Stochastic splicing has also been suggested by Wan et al. who observed large kinetic variation in intron removal for single introns in single cells (Wan et al., 2021). The indicated role of transcription factor concentrations (here, Eaf), suggests the need for a mechanism for local concentration buffering, e.g. by phase separation driven hub formation.

We provide initial measurements of splicing delay over developmental time, and we encourage their application in mathematical models; For example, to determine the relationship of parameters like Eaf concentration, nuclear volume increase and splicing delay over time. Since Cdk9 activity has been shown to affect splicing, it will be interesting to test if Eaf overexpression or depletion changes splicing delay rates.

Our previous finding of clustered histone modifications H3K27me3 and H3K4me3 marking pause and active transcription respectively as well as H3K36me3, which has been associated with splicing, along Y-loops is consistent with our model and will be another link to be explored in the future (Koestler et al., 2024; Shenasa and Bentley, 2023).

We have also addressed the question of how tightly the nRNA is coupled to the moving polymerase. The fact that most introns are spliced out co-transcriptionally means that the splice sites must be brought together, either in a transcription factory like arrangement with a stationary RPol and by looping of the DNA, or by carrying the upstream exon transcript along the intron coding DNA. The observed linkage of the 5’ splice site to the RPol by the U1snRNP likely ensures that the splice sites can readily be joined and suggests a rather tight coupling of nRNA and RPol (Beyer and Osheim, 1988; Cramer, 2019; Zhang et al., 2021). The huge introns of the Y-loop genes present an exaggerated example of a problem common to all splicing: how do the 5’ and 3’ splice sites find each other? In “regular” genes with short introns imaging has limited scope to provide insight as the short length of the introns forces the 5’ splice site to be close to the transcribing polymerase. However, the huge introns of the Y-loops mean that, in principle, the 5’ splice site could move several µm away from the site of transcription. This is not what we find and our imaging indicates that, during transcription of the large introns, the 5’ exon remains close to the transcribed chromatin fiber (Figs 1D, F and 2D). We suggest this is important for the process of bringing the 5’ and 3’ splice sites together. This proximity could potentially be achieved by transcript folding, by tethering to the polymerase (Zhang et al., 2021) or by trapping in the splicing hub.

Fingerhut et al. (Fingerhut et al., 2024) suggested that splicing must occur during transcription to prevent RNA entanglement. Our results indicate that also the intron containing nRNA is relatively compact and close to the RPol, which should counteract entanglement. The previously shown Christmas tree structure using electron microscopy, while beautifully illustrating transcription by staggered nRNA lengths, is a result of the preparation procedure by chromatin spreading rather than the *in-situ* arrangement (Miller and Beatty, 1969). We suggest that the decrease in transcription upon splicing interference is due to a halt in check point release after pausing, which may be more prominent in long genes. This would be consistent with Fingerhut et al. who observe a specificity of transcription attenuation upon splicing interference on long genes (Fingerhut et al., 2024).

Another factor in transcription/splicing kinetics to consider is the change of the Y-loop chromatin structure, which is assumed to decondense during SC maturation, but whose dynamics have not yet been analyzed in detail. For instance, initially transcription rate may be limited by condensed Y-loop chromatin, thus allowing splicing to take place, while the fully extended Y-loop in mature SCs allows continuous fast transcription leading to more delayed splicing.

Our observations also demonstrate the applicability and advantages of the *Drosophila* testis as a model system for splicing research: features such as co-transcriptional splicing, delayed splicing, transcription along the fiber and regulation of transcriptional activity are similar in a range of metazoans including humans. Most previous imaging studies relied on spot-like dimensions of a gene and its products, thus requiring indirect measurements and building on assumptions for the interpretation of results (Coté et al., 2024; Wan et al., 2021). Using the extended Y-loops, the sequential gene architecture, transcription and splicing activities are directly revealed by imaging of probes against different regions such as introns and exons.

The spatial and thereby also temporal separation of single events enables the kinetic analysis within single nuclei. Comparison between cells within the developing tissue informs about the influence of naturally occurring external factors versus hard wired regulation.

Determining the factors that regulate transcription pause and RPol speed will be central to understanding coordination of splicing and transcription and associated diseases, especially during cellular processes where conditions such as nuclear volume and transcription factor concentrations change (Northcott and Weaver, 2018; Roy, 1997).

## Methods

### Testes fixation

The procedure was based on (Raich et al., 2020). *Drosophila* melanogaster w1118 or CPTI-000560 (Venus-YFP::Eaf (Lowe et al., 2014)) testes from larvae or 0-2 d old males were dissected into Phosphate buffered saline (PBS) and fixed in a 1.5 ml microcentrifuge tube containing 300 µl 4 % Formaldehyde (FA, Sigma), 1 % Triton X-100 in PBS and 600 µl Heptane. The tube was shaken by hand for 30 s and fixation was continued on a rocker at room temperature for 30 min. Phases were allowed to separate, the solutions discarded, and the testes washed 5x with 0.01 % Triton X-100 in PBS (PTX0.01).

### Stage determination during spermatogenesis

This was performed on fixed and DAPI stained whole-mount testes as shown in Fig. 4A. Stages were determined according to Cenci et al. (Cenci et al., 1994).

### HCR-RNA-FISH

The procedure was done as in Sharrock et al. (Sharrock et al., 2022). The probe sets were designed and produced by Molecular Instruments Inc. (USA) according to the target sequences (see supplemental material SuppText_RNAFISHsequences.txt). Briefly, fixed testes were incubated overnight at 37°C with hybridization solution containing the HCR probe sets and oligonucleotide probe. This was followed by washing and amplification with hairpins overnight. Testes were washed, stained with DAPI (0.5 µg/ml) and mounted in glycerol.

HCR-FISH probe target region lengths:

*kl-5* intron 1 3’ region: 3438 bp

*kl-5* exon block 2-9 (without introns): total 2167 bp

*kl-5* intron 9 5’ region: 1710 bp

*kl-5* intron 9 central region: 13141 bp

*kl-5* intron 9 3’ region: 7165 bp

*kl-5* exons10-12 plus flanking sequences including introns: 4980 bp

6xAAGAC_repeat_ATTO565 oligonucleotide probe: 30 nt

### Immunofluorescence staining

Fixed testes were permeabilized and blocked with 0.5 % Triton X-100, 1% WBR (Western Blot reagent, Roche) in PBS o/n at 4°C, then incubated with primary antibodies (in PBS/1%WBR) for 2h, washed 2x 20 min with PT0.1 (0.1 % Tween 20 in PBS) and kept overnight at 4°C. This was followed by incubation with secondary antibody for 1h at room temperature. After washing 3x 20 min with PT0.1 at RT, testes were postfixed (3.7% FA / PBS 10 min) washed with PTX0.01 and mounted with glycerol.

Primary antibodies were: rabbit anti-RPol-PSer2 (ab238146, Abcam), 1:500; rabbit anti-RPol-PSer5 (ab76292, Abcam, 1:500), mouse anti-Sm antigen (United States Biological S1014-29BX; 1:200). Secondary antibodies were from Invitrogen and used at 1:1000; goat anti-mouse Ig-Alexa Fluor 647 (A-21235), goat anti-rabbit Ig Alexa Fluor 488 (A-11008) and goat anti-rabbit Ig Alexa Fluor 568 (A-11011).

### Microscopy and image processing

Fixed HCR-RNA-FISH specimens (Figs. 1–3) were imaged as z-stacks on a Zeiss LSM 900 inverted microscope (Axio Observer) with an Airyscan 2 detector and a Plan-Apochromat 63×/1.40 oil-immersion objective. The four fluorescence channels were acquired in separate tracks with 405, 488, 561 and 640 nm excitation, bidirectional scanning and twofold or fourfold line averaging. Recorded emission detection windows for these excitation tracks were 400–605, 490–555, 545–620 and 450–700 nm, respectively; in Fig. 1B/C they were 400–565, 490–565, 555–620 and 620–700 nm. The processed stacks had 0.0353 × 0.0353 µm pixels and 0.14 µm z-spacing. Fig. 1B/C combined HCR probes for exons 2–9 (Alexa Fluor 488) and exons 10–12 (Alexa Fluor 647) with an (AAGAC)₆ repeat oligonucleotide (ATTO 565). Fig. 1D and Figs. 2&3 combined probes for the 3′ region of intron 1 (Alexa Fluor 647), exons 2–9 (Alexa Fluor 488) and the 5′ region of intron 9 (Alexa Fluor 546). Fig. 1E combined probes for exons 2–9 (Alexa Fluor 488), central intron 9 (Alexa Fluor 647) and the 3′ region of intron 9 (Alexa Fluor 546); Fig. 1F combined probes for the 5′ region of intron 9 (Alexa Fluor 488), the 3′ region of intron 9 (Alexa Fluor 546) and exons 10–12 (Alexa Fluor 647). All FISH specimens were counterstained with DAPI. The images in Fig. 1B/C and 1E underwent ZEN Airyscan 3D processing; the Fig. 1B/C stack additionally underwent stack correction. The Fig. 1D, 1F and Figs. 2&3 stacks underwent Airyscan joint deconvolution. Fig. 1B/C, 2 testes, Fig. 1D, 2 testes, Fig. 1E, 1 testis, Fig. 1F, 1 testis, Fig. 2&3, 2 testes.

Immunofluorescence specimens (Figs. 5&6) were imaged with the Zeiss Airyscan system. RNA polymerase II phospho-Ser2 (Fig. 5) or phospho-Ser5 (Fig. 6) was detected in the Alexa Fluor 647 channel; anti-Smith was detected in the Alexa Fluor 568 channel, endogenous Eaf-Venus-YFP by direct fluorescence, and DNA by DAPI staining. In the Fig. 5&6 acquisitions, recorded emission windows for 405, 488, 561 and 640 nm excitation were 400–530, 495–580, 575–620 and 450–700 nm, respectively. Four testes were imaged per condition.

Fixed whole-mount testes labelled for Eaf-Venus-YFP, EU–Alexa Fluor 488 and DAPI (Fig. 4A/F) were imaged on a Leica TCS SP8 confocal microscope with an HC PL APO CS2 40×/1.30 oil-immersion objective. Z-stacks were acquired with bidirectional scanning, sequential fluorescence acquisition and fourfold to sixteenfold line averaging; the recorded xy pixel sizes ranged from 0.06 to 0.14 µm and z-steps from 0.35 to 0.7 µm. Spectral detector windows were 410–501, 519–610 and 638–760 nm. Six testes were imaged.

Eaf-Venus-YFP was imaged live in freshly dissected larval whole-mount testes in medium(Sigma Schneider’s +10% FBS (Biosera, HI)) at 25°C in an imaging chamber with a #1.5H glass coverslip (Ibidi) as time-lapse z-stacks (Fig. 4C/D; Video 2), at one z-stack per 31.52 s (Fig. 4C), 20.00 s (Fig. 4D) or 31.42 s (Video 2). 18 nuclei in 3 testes were imaged.

For the 1,6-hexanediol assay (Fig. 4E), two independent testes were imaged on the Zeiss system in confocal mode, at one z-stack per 120.35 s or 150.64 s. After 5–10 min of baseline imaging in 500 µl medium, testes were exposed to 5% 1,6-hexanediol (Sigma-Aldrich) in medium for 14–42 min. The solution was then replaced with medium alone and recovery was imaged for 30 min.

Zeiss images were processed in ZEN (blue edition); Leica images were acquired in LAS X. Images were inspected and prepared using Fiji/ImageJ.

Image metadata for this section were summarized using GPT6-Sol (OpenAI).

### EU labeling

Dissected testes were resuspended in medium (Sigma Schneider’s +10% FBS (Biosera, HI)) containing 1 mM EU (5-ethynyl uridine; Molecular Probes C10330) for 20 min at 25°C and fixed as above. For the Click reaction the manufacturer’s manual was followed (Molecular Probes C10330) using Alexa Fluor 647 Azide (Invitrogen A10277). Briefly, testes were incubated with Click-iT reaction cocktail for 45 min and washed with reaction rinse buffer.

### Image analysis

Splicing delay measurements within single nuclei: We estimated the splicing delay of intron 1 by measuring the distance along a loop covered by FISH signal in Fiji. The obtained values must be considered as minimal estimates for several reasons: measurements were performed on maximum intensity projections; the orientations of the loops differ considerably; the full path cannot be unambiguously identified.

Splicing delay measurements along the testis: Nuclei were cropped out as z-stacks, followed by z-projection (sum), an ROI drawn around manually (z-projected) Y-loop intron/exon signals to measure their integrated intensities. These values were normalized to the captured maxima and plotted over the distance of the center of each nucleus to the point of highest maturity. The point of highest spermatocyte maturity was defined by the most proximal and central point within the field of view (capturing S4 to S6 primary spermatocytes) and was guided by transcript probe signal and the density of cytoplasmic exon signal containing particles.

The overlap of YFP-Eaf and immunostained RPol-PSer2 or -PSer5 was determined following segmentation using Arivis Vision 4D.

Eaf intensity plots along testes were performed on maximum intensity projections of z-stacks of YFP-Eaf expressing testes by drawing a line along testes. Line width was adjusted individually for each measurement. Intensity was measured with the plot profile tool in Fiji/ImageJ and normalized to the maximum and minimum values. Trendlines were added in Excel (Microsoft) with a polynomial 2.

Statistics were performed in Excel (Microscoft).

For image presentation, 3D rendering and video production Fiji/ImageJ (Schindelin et al., 2012), Arivis Vision 4D (Zeiss), Aivia (Leica), DaVinci Resolve (Blackmagic Design) and Shutter Encoder were used.

## Acknowledgements

We thank the Cambridge Advanced Imaging Centre (CAIC) for help with microscopes.

## Supplementary files

Supplementary files can be downloaded from Zenodo.org: 10.5281/zenodo.24094594

Video 1: corresponds to Fig. 1B. Testis overview with FISH of intronic repeats, exons 2-9 and exons 10-12.

Video 2: Corresponds to Fig. 4CD, but showing a different testis, showing Eaf speckle dynamics in whole-mount testes. Frame interval = 31.42 s.

Video 3: Shows the nucleus of Fig. 4C, with a “breathing” Eaf speckle. Frame interval = 31.52 s.

Video 4: Corresponds to Fig. 4D, where an Eaf speckle moves and fuses with another one at the periphery.

Video 5: Corresponds to Fig. 4E, Hexanediol treatment, leading to dissolution of Eaf speckles and reformation after washout.

Videos were prepared with ImageJ/Fiji, Aivia (Leica) and DaVinci Resolve (Blackmagic Design).

Supplementary Figure: corresponds to Fig. 4B, Eaf intensities along testes. It shows the image data, measurement regions and raw intensity data.

Supplementary data: provides the measurements to Fig. 2A/B, 2C and 5B.

Supplementary text: provides the RNA-FISH target sequences.

